# ExoRNAi exposes contrasting roles for sugar exudation in host-finding by plant pathogens

**DOI:** 10.1101/015545

**Authors:** Neil D Warnock, Leonie Wilson, Juan V Canet-Perez, Thomas Fleming, Colin C Fleming, Aaron G Maule, Johnathan J Dalzell

## Abstract

Plant parasitic nematodes (PPN) locate host plants by following concentration gradients of root exudate chemicals in the soil. We present a simple method for RNAi-induced knockdown of genes in tomato seedling roots, facilitating the study of root exudate composition, and PPN responses. Knockdown of sugar transporter genes, *stp1* and *stp2* in tomato seedlings triggers corresponding reductions of glucose and fructose, but not xylose, in collected root exudate. This corresponds directly with reduced infectivity and stylet thrusting of the promiscuous PPN *Meloidogyne incognita,* however we observe no impact on the infectivity or stylet thrusting of the selective Solanaceae PPN *Globodera pallida.* This approach can underpin future efforts to understand the early stages of plant-pathogen interactions in tomato, and potentially other crop plants.

---

RNA interference (RNAi) is widely used for the analysis of plant gene function, primarily through the transgenic production of dsRNA constructs *in planta,* and secondarily through Virus-Induced Gene Silencing (VIGS) (Watson *et al.,* 2005). Publications from the Wolniak lab have shown that exogenous dsRNA can silence genes of the water fern *Marsilea vestita* (Klink and Wolniak, 2001), and crude lysate from *Escherichia coli* expressing virus-specific dsRNA have also been used to protect plants from viral pathology (Tenllado *et al.,* 2003). Here we present a similar approach to triggering RNAi in tomato seedlings, which we term exogenous (exo)RNAi. In this approach, aqueous dsRNA is delivered exogenously to tomato seedlings.

Plant root exudate comprises a complex mixture of compounds including volatile and soluble chemicals which may derive from intact or damaged root cells, or sloughed-off root border cells (Dakora and Phillips, 2002). It has been estimated that 11% of photosynthetically-assimilated carbon is released as root exudate (Jones *et al.,* 2009). The monosaccharides glucose, fructose and xylose represent the major sugar component of tomato root exudates (Kamilova *et al.,* 2006). Plant parasitic nematodes (PPNs) are responsible for an estimated 12.3% loss in crop production globally each year (Sasser and Freckman, 1987), and are attracted to host plants by components of plant root exudate. Here we assess the chemosensory response of the root knot nematode, *Meloidogyne incognita* (a promiscuous pathogen of flowering plants), and the potato cyst nematode, *Globodera pallida* (a selective pathogen of Solanaceae plants) to each of the three major monosaccharide sugars of tomato plant root exudate, and the efficacy of exoRNAi against *stp1* and *stp2,* known transporters of monosaccharide sugars in tomato (Gear *et al.,* 2000).

*Meloidogyne incognita* infective stage juveniles were attracted to glucose (CI: 0.33 ±0.07; P<0.001) and fructose (CI: 0.39 ±0.09; P<0.001), but not xylose (CI: 0.04 ±0.09; P>0.05) as compared to control treated worms (Fig 1A). Glucose (125.1% ±5.5; P<0.001) and fructose (124.8% ±5.4; P<0.001) also triggered an elevated level of serotonin-triggered stylet thrusting in treated juveniles; xylose failed to trigger any significant response (99.36% ±10.87; P>0.05) when compared to control treatments (Fig 1B). *Globodera pallida* infective stage juveniles were mildly repelled by glucose (CI: -0.23 ±0.09; P>0.05), and did not respond to fructose (CI: 0.15 ±0.08; P>0.05), or xylose (CI: -0.19 ±0.09; P>0.05) as compared to control treated worms (Fig 1C). Glucose (118.6% ±9.7; P>0.05), fructose (107.2% ±7.3; P>0.05), or xylose (119.6% ±8.6; P>0.05) had no significant impact on the frequency of serotonin-triggered stylet thrusting in *G. pallida* infective juveniles when compared to control treatments (Fig 1D).

**Figure 1.**
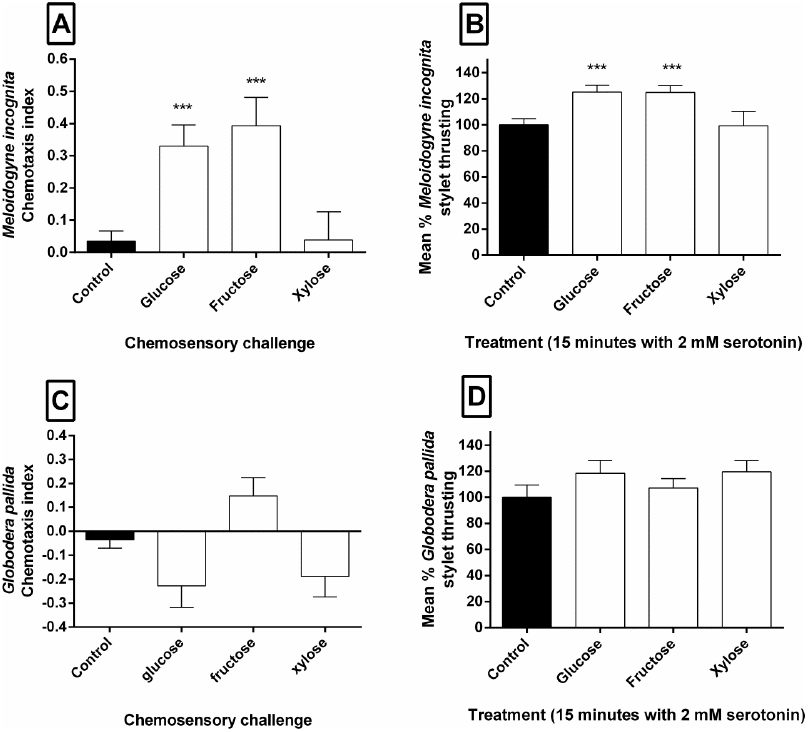
Glucose and fructose stimulate different chemotaxis and stylet thrusting responses in *M. incognita* and *G. pallida.* **(A)** Chemosensory response (chemosensory index) of *M. incognita* infective juveniles to glucose, fructose, xylose and control (water) assay challenge. Each data point represents the mean (±SEM) of 10 assays of 100 infective juveniles each. **(B)** Mean percentage (±SEM) stylet thrusting of glucose, fructose and xylose treated *M. incognita* infective stage juveniles (n=100) relative to control (2 mM serotonin in water). **(C)** Chemosensory response of *G. pallida* infective juveniles to glucose, fructose, xylose and control (water) assay challenge. **(D)** Mean percentage (±SEM) stylet thrusting of glucose, fructose and xylose treated *G. pallida* infective stage juveniles (n=100) relative to control (2 mM serotonin in water). An agar slurry (0.25% agar, pH 7) was used to flood Petri dishes for chemosensory assays. Specifically, 3 ml of agar slurry was poured to provide the medium through which the infective stage juveniles could move. Sugar plugs were prepared by dissolving 50 mM of the relevant sugar (glucose / fructose / xylose) in 0.25% agar and allowed to set. Plugs were picked with a Pasteur pipette which had been cut half way down the pipette barrel, and placed onto one side of a Petri dish, with a negative plug (water instead of 50 mM sugar) on the other. 100 *M. incognita* or *G. pallida* infective stage juveniles were suspended in 5 μl of water, and spotted onto the centre point of each dish. A Petri dish lid was marked with two parallel vertical lines 0.5 cm either side of the centre point forming a 1 cm ‘dead zone’ that ran vertically along the lid. Assay plates were set onto the lid for scoring of nematode positions following a two hour assay period. Only nematodes outside the dead zone were counted. The distribution of *M. incognita* infective stage juveniles was used to generate the chemotaxis index (Hart, 2006) for each assay plate which formed one replicate. For the stylet thrusting assay, 100 *M. incognita* or *G. pallida* infective stage juveniles were suspended in 20 μl of water (autoclaved and adjusted to pH 7) containing 2 mM serotonin and 50 mM of glucose, fructose or xylose (Sigma-Aldrich). Worms were incubated in this solution for 15 minutes, pipetted onto a glass slide with a coverslip, and stylet thrusts were counted in randomly selected infective stage juveniles for 1 minute each. Control treatments were expressed as a percentage, including technical variation, and experimental treatments were normalised to control percentages across individual experiments and days. Chemosensory and stylet thrusting results were analysed by One-way ANOVA and Tukey’s Honestly Significant Difference test using Graphpad Prism 6. Probabilities of less than 5% (P < 0.05) were deemed statistically significant *, P<0.05; **, P<0.01; ***, P<0.001.

Treatment of tomato seedlings with *stp1* dsRNA triggered a significant reduction in *stp1* transcript abundance (0.17 ±0.05; P<0.001), yet had no impact on *stp2* abundance (1.037 ±0.13; P>0.05) relative to neomycin transferase (neo) dsRNA treatment. Likewise, *stp2* dsRNA induced significant reductions in *stp2* transcript abundance (0.21 ±0.06; P<0.001), but not *stp1* (0.94 ±0.05; P>0.05) relative to *neo* dsRNA treatments (Fig 2A). Corresponding reductions in glucose and fructose exudate concentration were observed for both *stp1* (5.10 μg/ml ±1.31; P<0.01 and 3.14 μg/ml ±0.92; P<0.01, respectively) and *stp2* (4.90 μg/ml ±1.45; P<0.01 and 10.90 μg/ml ±1.07; P<0.05, respectively) dsRNA treated seedlings. No significant changes in xylose exudate concentration were observed across treatment groups (Fig 2B-D).

**Figure 2.**
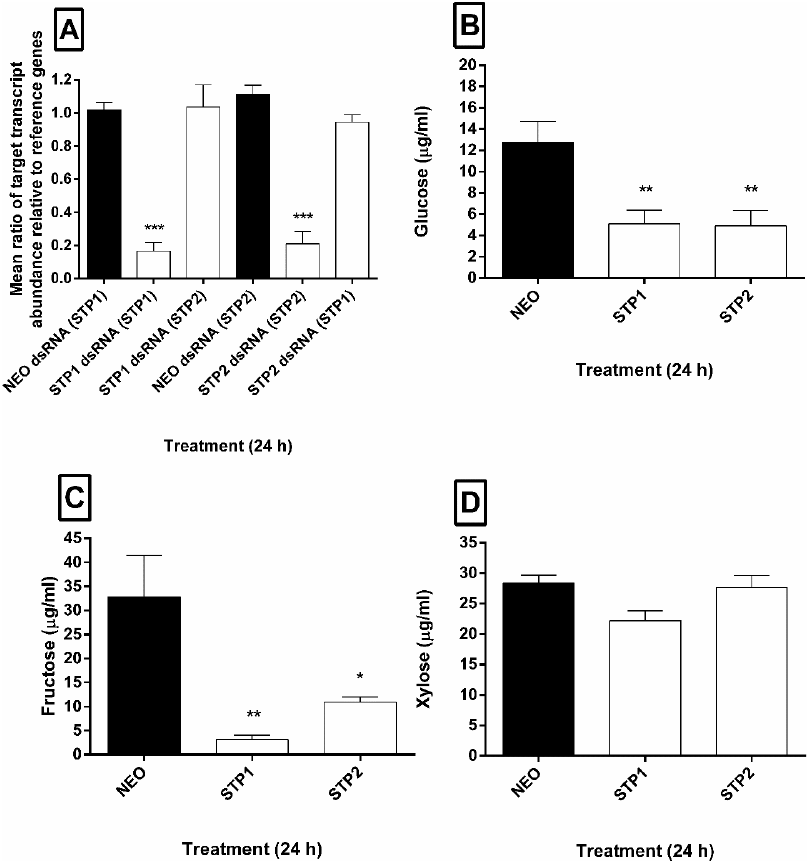
ExoRNAi induces target-specific knockdown of tomato sugar transporter genes; alters root exudate composition. **(A)** Mean ratio of target transcript (in parentheses) abundance relative to three endogenous reference genes. Each data point represents the mean (±SEM) of three replicates of five seedlings each. Forward and reverse primers including 5’ T7-recognition sites were used to generate specific amplicons for dsRNA synthesis to EST-supported fragments of *stp1* (Solyc02g079220.2), and *stp2* (Solyc09g075820.2) (Reuscher *et al.,* 2014). Primers for dsRNA synthesis were as follows: (Neomycin Phosphotransferase, *neo*F 5’ -GGTGGAGAGGCTATTCGGCT-3’, neoFT7 5’-TAATACGACTCACWAGGGGTGGAGAGGCTATTCGGCT -3’, neoR 5’- CCTTCCCGCTTCAGTGACAA-3’, neoRT7 5’-T AAT ACGACT CACT AT AGGCCTT CCCG CTT CAGT GACAA -3’); (Sugar Transporter 1, *stp1*F 5’- CTGCTGTGATCACTGGTGGA-3’, stp1FT7 5’-T AAT ACGACT CACT AT AGGCT GCT GT GAT CACT GGTGGA -3’, *stp1R* 5’-ATT CCCCTGGAGTTCCATTT-3’, *stp1* RT7 5’- T AAT ACGACT CACT AT AGGATT CCCCTG GAGTTCCATTT -3’); (Sugar Transporter 2, *stp2*F 5’- ACGTTCTCTCCACCGTTGTC -3’, *stp2*FT7 5’-TAATACGACTCACTATAGGACGTTCTCTCCACCGTTGTC -3’, *stp2*R 5’- CTACGAAGATTCCCCAACCA-3’, *stp2*RT7 5’-TAATACGACTCACTATAGGCTACGAAGATTCCCCAACCA-3’); PCR products were assessed by gel electrophoresis, and cleaned using the Chargeswitch PCR clean-up kit (Life Technologies). dsRNA was synthesised using the T7 RiboMAX™ Express Large Scale RNA Production System (Promega), and quantified by Nanodrop 1000 spectrophotometer. Tomato cv. Moneymaker seeds (Suttons) were sterilised by 30 minute treatment in dilute bleach, followed by five, 15 minute washes in 1 ml deionised water. Seeds were germinated on 0.5X MS salts, 0.6% agar plates at 23°C, and taken for exoRNAi treatment on the first day post radicle emergence. Ten seedlings were used per well of a 24-well plate (SPL Lifesciences), and incubated with 300 μl of 10 ng/μl dsRNA solution for 24h at 23°C, in darkness. Five seedlings were snap frozen in liquid nitrogen per biological replicate, and total RNA isolated using Trizol reagent. Total RNA was treated with the Turbo DNase free kit (Life Technologies), and cDNA was synthesised using the High-capacity RNA-to-cDNA kit (Applied Biosciences) according to manufacturer’s instructions using the maximum input concentration of RNA. Three biological replicates were performed for each treatment. Quantitative RT-PCR primers were as follows: (Sugar Transporter 1, qstp1F 5’-ATGTTGCTGGATTCGCTTGGTC-3’, qstp1R 5’- TGTGCAGCTGATCGAATTTCCAG-3’); (Sugar Transporter 2, qstp2F 5’-ATTATGGCTGCTACCGGAGGTC-3’, qstp2R 5’-TGTAACACCACCAGAAACTCCAAC-3’); (Elongation Factor, *qefα*F 5’-TACTGGTGGTTTTGAAGCTG-3’, qefaR 5’- AACTTCCTTCACGATTTCATCATA-3’); (SAND protein family, qsandF 5’- TTGCTTGGAGGAACAGACG-3’, qsandR 5’- GCAAACAGAACCCCTGAATC-3’); (Sugar Transporter 41, *qstp41*F 5’- ATGGAGTTTTTGAGTCTTCTGC -3’, *qstp41R* 5’-GCTGCGTTTCTGGCTTAGG -3’) (Dekkers *et al.,* 2012). Primer sets to be used for qPCR were optimised for working concentration, annealing temperature and analysed by dissociation curve for contamination or nonspecific amplification by primer–dimer as standard. Each individual reaction comprised 5 μl Faststart SYBR Green mastermix (Roche Applied Science), 1 μl each of the forward and reverse primers (10 mM), 1 μl water, 2 μl cDNA. PCR reactions were conducted in triplicate for each individual cDNA using a Rotorgene Q thermal cycler under the following conditions: [95°C × 10 min, 40 × (95°C × 20s, 60°C × 20s, 72°C × 20s) 72°C × 10 min]. The PCR efficiency of each specific amplicon was calculated using the Rotorgene Q software, and quantification of each target amplicon calculated by an augmented comparative Ct method (Pfaffl, 2001), relative to the geometric mean of three endogenous reference genes (Vandesompele *et al.,* 2002). Ratio-changes in transcript abundance were calculated relative to control dsRNA treated seedlings in each case Exudate concentration of **(B)** glucose, **(C)** fructose and **(D)** xylose across *neo* (double stranded [ds]RNA control), *stp1* and *stp2* dsRNA treated tomato seedlings. The exudate solution was collected by pipette and transferred to a hydrophobically-lined microcentrifuge tube (Anachem) prior to quantification. The sugars were quantified colorimetrically at 340 nm using Glucose (HK), and Fructose assay kits from Sigma-Aldrich, and the Xylose assay kit from Megazyme as per manufacturer’s instructions. Each data point represents the mean (±SEM) of three replicates of ten seedlings each. Data were analysed by ANOVA and Tukey’s Honestly Significant Difference test using Graphpad Prism 6. Probabilities of less than 5% (P < 0.05) were deemed statistically significant *, P<0.05; **, P<0.01; ***, P<0.001.

Root exudates collected from tomato seedlings which had been treated with either *stp1* or *stp2* dsRNA were less capable of stimulating stylet thrusting in *M. incognita* relative to exudates collected from control dsRNA treated seedlings (13.92 ±5.10%, P<0.001; and 17.53 ±8.12%, P<0.001, respectively. Fig 3A). No significant difference in stylet thrusting frequency was observed for *G. pallida* juveniles when exposed to root exudates from *stp1* or *stp2* dsRNA-treated seedlings, relative to control treated seedlings (108.2 ±38.87%, P>0.05; and 77.34 ±30.84%, P>0.05, respectively) (Fig 3B).

When exoRNAi-treated seedlings were challenged by *M. incognita* infection assay, significant reductions in percentage infection levels relative to control (neo) dsRNA treatment were observed for both *stp1* (14.15% ±4.77; P<0.01) and *stp2* (27.08% ±7.32; P<0.05) dsRNA treatments (Fig 3C). Knockdown of *stp1* (14.15% ±4.77; P>0.05) or *stp2* (14.15% ±4.77; P>0.05) did not significantly reduce the percentage infection levels of *G. pallida* relative to *neo* dsRNA treatment (Fig 3D).

**Figure 3.**
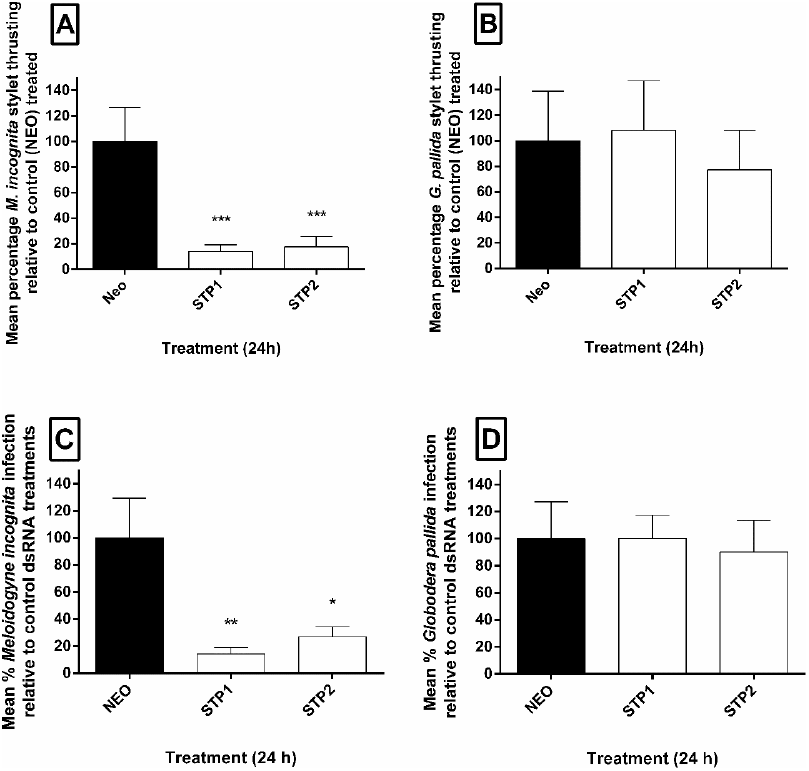
ExoRNAi of tomato seedling sugar transporters differentially alters plant nematode infection and activation. (A) Mean percentage (±SEM) stylet thrusting of *M. incognita* and (B) *G. pallida* infective stage juveniles in response to collected seedling exudates (n=100), relative to control *(neo* dsRNA). Root exudate was collected and quantified as described in Fig. 2. Nematodes were exposed to 50μl of dsRNA treated seedling exudate for 30 minutes, thrusts were counted in randomly selected infective stage juveniles for 1 minute each. (C) Mean percentage *M. incognita* infection levels of *stp1* and *stp2* dsRNA treated tomato seedlings normalised to control (neo) dsRNA treated seedlings. (D) Mean percentage *G. pallida* infection levels of *stp1* and *stp2* dsRNA treated tomato seedlings normalised to control (neo) dsRNA treated seedlings. Agar slurry was prepared by autoclaving a 0.55% agar solution which had been autoclaved and adjusted to pH 7. The agar was agitated for six hours at room temperature, until it had a smooth consistency. 500 *M. incognita* or *G. pallida* infective stage juveniles were added to each well of a 6 well plate (SPL Lifesciences) with one exoRNAi treated seedling embedded within 3 ml of agar slurry. Plates were sealed with parafilm, covered above and below with a sheet of tin foil and incubated for 24 hours at 23°C. Seedlings were subsequently removed from the slurry, gently washed several times by immersion in deionised water, and stained using acid fuschin (Bybd *et al.,* 1983). The number of invading PPN juveniles was counted for each seedling using a light microscope. Control treatments were expressed as a percentage, including technical variation, and experimental treatments were normalised to control percentages. Each data point represents the mean (±SEM) of ten seedlings challenged with 500 infective stage juveniles each. *, P<0.05; **, P<0.01; ***, P<0.001.

These data demonstrate that the exogenous application of aqueous dsRNA onto tomato seedlings is sufficient to trigger specific gene knockdown. However, we found that different experimental populations of tomato seedlings could display wide variation in the expression of both sugar transporter genes, and reference genes which resulted in high standard error of mean (SEM) values. This made it difficult to resolve gene knockdown levels for an isolated number of experiments. This may be due to variation in the susceptibility of tomato seedlings to exoRNAi, as has been observed for Tobacco Rattle Virus (TRV) VIGS approaches in tomato (Liu *et al.,* 2002), or it could indicate that larger replicates of seedlings are required to consistently resolve gene expression data post exoRNAi. It should also be noted that attempts to silence phytoene desaturase in order to observe a bleaching phenotype in the cotyledons were unsuccessful (data not shown). This may indicate that only genes expressed in the tomato root are susceptible to this approach.

It is well established that plant root exudates mediate both positive and negative interactions with commensal and pathogenic microbes (Badri *et al.,* 2009), insects (Walker *et al.,* 2003), and other plants (Bais *et al.,* 2006). Plant parasitic nematodes also respond to plant root exudates (Teillet *et al.,* 2013). The present study aimed to probe the involvement of monosaccharide sugars of tomato root exudate for involvement in the attraction and activation of parasitic behaviours in the promiscuous root knot nematode *M. incognita,* and the host-selective potato cyst nematode *G. pallida.* STP1 and STP2 are known transporters of monosaccharide sugars (Gear *et al.,* 2000), and our data demonstrate that both play a role in regulating the level of glucose and fructose (but not xylose) exudation from tomato seedling roots. exoRNAi knockdown of each transporter significantly reduced the amount of glucose and fructose secreted from plant roots, which corresponded with a decrease in *M. incognita* infectivity, but not *G. pallida* infectivity. These results suggest that glucose and fructose are important chemical cues which infective stage *M. incognita* use to find host plants. These data indicate that glucose and fructose trigger host-finding and stylet thrusting in promiscuous PPNs, as opposed to host-specific PPNs, an observation which is consistent with the ubiquitous nature of monosaccharide sugars in plant root exudates (Kamilova *et al.,* 2006). The demonstration that STP1 and STP2 are specifically involved in the exudation of both monosaccharides from tomato roots is an important finding which can underpin future efforts to study the link between plant root transporters, and chemical constituents of root exudates.

## Acknowledgements

This work was supported financially by Queen’s University Belfast. Dalzell is supported by an Early Career Fellowship from The Leverhulme Trust. Warnock was supported by a Gates Foundation Grand Challenges Grant, Wilson was supported by a EUPHRESCO fellowship, Canet-Perez was supported by an Invest Northern Ireland Proof-of-Concept award and Fleming by a Department of Agriculture and Rural Development studentship award.

